# Suboptimal intermediates underlie evolution of the Bicoid homeodomain

**DOI:** 10.1101/2020.08.20.257824

**Authors:** Pinar Onal, Himari Imaya Gunasinghe, Kristaley Yui Umezawa, Michael Zheng, Jia Ling, Leen Azeez, Anecine Dalmeus, Tasmima Tazin, Stephen Small

## Abstract

Changes in regulatory networks generate materials for evolution to create phenotypic diversity. For transcription networks, multiple studies have shown that alterations in binding sites of cis-regulatory elements correlate well with the gain or loss of specific features of the body plan. Less is known about alterations in the amino acid sequences of the transcription factors (TFs) that bind these elements. Here we study the evolution of Bicoid (Bcd), a homeodomain (HD) protein that is critical for anterior embryo patterning in *Drosophila*. The ancestor of Bcd (AncBcd) emerged after a duplication of a Zerknullt (Zen)-like ancestral protein (AncZB) in a suborder of flies. AncBcd diverged from AncZB, gaining novel transcriptional and translational activities. We focus on the evolution of the HD of AncBcd, which binds DNA and RNA, and is comprised of four subdomains: an N-terminal arm (NT) and three helices; H1, H2, and Recognition Helix (RH). Using chimeras of subdomains and gene rescue assays in *Drosophila*, we show that robust patterning activity of the Bcd HD (high frequency rescue to adulthood) is achieved only when amino acid substitutions in three separate subdomains (NT, H1, and RH) are combined. Other combinations of subdomains also yield full rescue, but with lower penetrance, suggesting alternative suboptimal activities. Our results suggest a multi-step pathway for the evolution of the Bcd HD that involved intermediate HD sequences with suboptimal activities, which constrained and enabled further evolutionary changes. They also demonstrate critical epistatic forces that contribute to the robust function of a DNA-binding domain.

## Introduction

The main components of transcription networks are transcription factors (TFs) and the cis-regulatory elements of target genes that contain TF-binding sites [1]. During evolution, DNA sequence changes in either component can alter network topology, affect gene expression patterns, and ultimately induce functional changes that are selected by evolutionary pressures. A sequence change in a cis regulatory element might affect the expression of a single gene, and it is thought that the evolution of body plan diversity is mainly driven by the accumulation of many such incremental changes [2]–[4]. In contrast, an amino acid change that alters the DNA-binding activity of a TF would alter the expression of many target genes and cause diverse and pleiotropic effects, which might be less compatible with survival. Despite this bias, several studies in plants and animals suggest that changes in TF sequences are critical for establishing variation during evolution [5]–[7].

One issue with the TF evolution hypothesis is that changes in TF function that generate new functions might interfere with critical roles normally played by the TF. However, this issue can be mitigated by gene duplication events, which provide extra genetic material for the evolution of novel or modified functions [8]–[12]. For example, multiple duplications in the Hox locus, followed by diversification of individual genes, were critical for establishing divergent body plans throughout the metazoa [13]–[19].

Here, we study the evolution of Bicoid (Bcd), a homeodomain (HD)-containing transcription factor that is critical for patterning anterior regions of the *Drosophila* embryo. The ancestor of *Drosophila **b**cd* gene (*ancbcd*) emerged 150 MYA in Cyclorrhaphan flies (a suborder in flies) after a duplication of an ancestral gene (*anczb*), which also gave rise to the ancestor of *bcd*’s sister gene *zerknullt (anczen)* [20]–[22]. In *Drosophila* and most other Cyclorrhaphan flies, *anczen* maintained an ancestral role in extraembryonic patterning, while *ancbcd* evolved rapidly. In addition to evolution in regulatory sequences that led to maternal expression and anterior localization of *bcd* mRNA, coding sequence changes completely altered the DNA-binding activities of AncBcd [23] and allowed it to bind to RNA [24], [25]. In the early embryo, Bcd protein is distributed in an anterior to posterior (AP) gradient [26] and is essential for transcriptionally activating more than 50 genes in unique temporal and spatial patterns along the AP axis [27]–[29]. Most Bcd target genes encode transcription factors, which regulate cross-regulate each other in a network that forms and positions seven head segments and three thoracic segments in the anterior half of the developing embryo [30]. Bcd also binds directly to the mRNA of the posterior determinant *caudal* (*cad*), and prevents its translation in anterior embryonic regions [31], [32]. Embryos lacking Bcd form no head or thoracic segments, and show variable defects in abdominal segmentation [33].

The Zen and Bcd proteins in *Drosophila* have completely different functions *in vivo*. Specifically, when expressed in a Bcd-like gradient in embryos lacking Bcd, Zen has no Bcd-like activity [34]. However, when the Bcd HD is swapped into the Zen protein, the chimeric ZenBcdHD partially rescues the morphological defects in Bcd-depleted embryos, and activates a subset of Bcd target genes [34]. These results indicate that the unique patterning activities of Bcd are determined in large part by its DNA- and RNA-binding preferences. They suggest further that amino acid substitutions in the AncZB HD were critical for the evolution of Bcd’s functions in anterior embryo patterning.

In a previous study, ancestral protein reconstruction (APR; [35]) was used to infer the amino acid sequences of the HDs that were present in AncZB and AncBcd. There are 31 high confidence differences between the two HDs, which are distributed among four HD subdomains: the N-terminal arm (NT) and three alpha helices [H1, H2, and the DNA recognition helix (RH)] (Figure 1A). When tested *in vivo*, a Bcd protein containing the AncZB HD failed to provide any Bcd-like activity, while an identical construct carrying the AncBcd HD completely rescued *bcd* deficient embryos to adulthood. [34], This study also tested the roles of two substitutions in the RH (q50>**K** and m54>**R**) because these had previously been shown to be critical for Bcd’s DNA-binding specificity [36], [37] and RNA-binding activities [31]. Substituting both the K50 and R54 amino acids into the AncZB resulted in the activation of a subset of Bcd target genes, but only partially rescued the morphological defects of embryos lacking Bcd, suggesting that other substitutions in the RH or in other subdomains of the HD were required for the evolution of AncBcd HD’s full transcriptional and post-transcriptional activities (Figure 1B).

**Figure 1:**
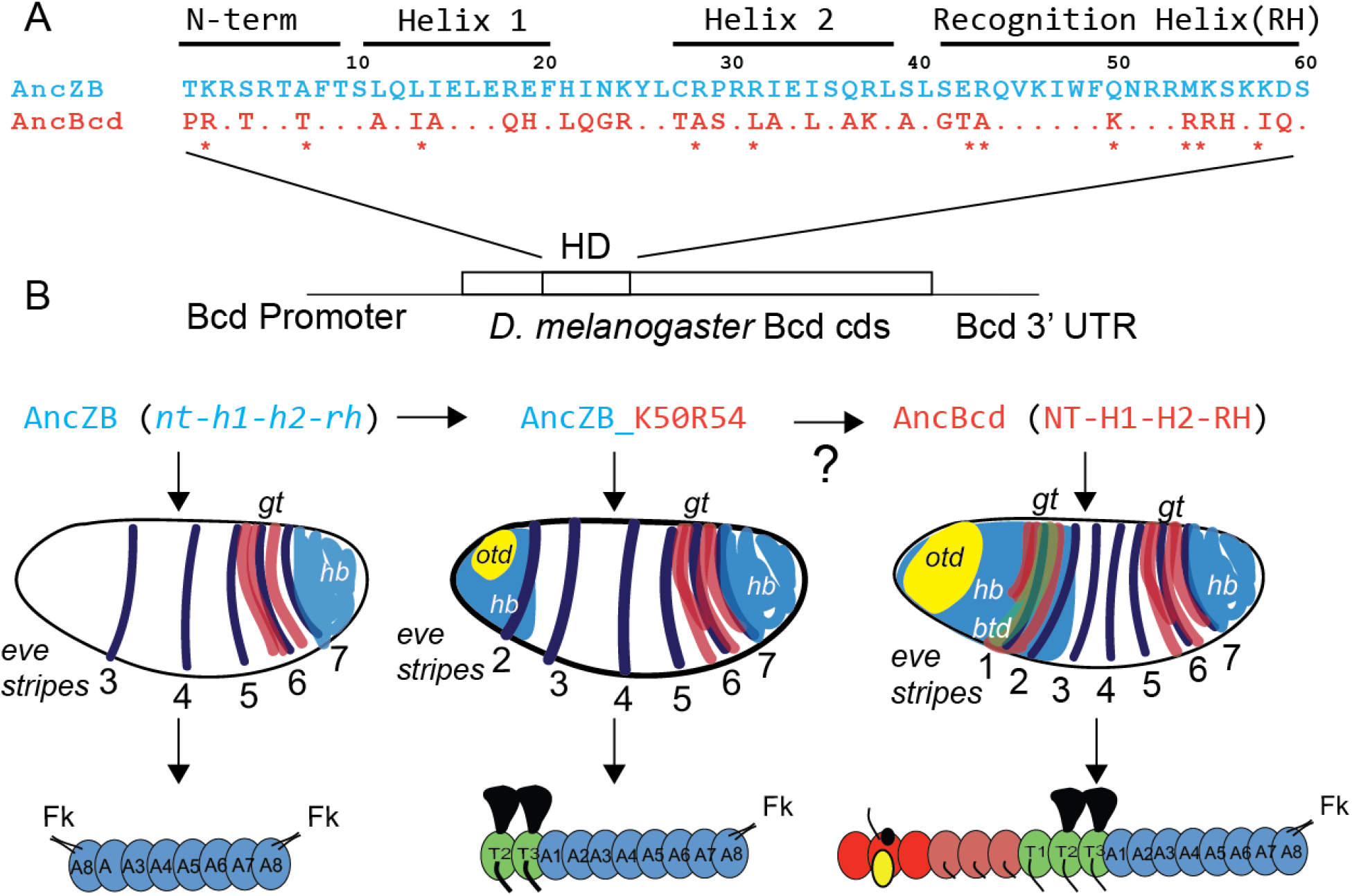
**A.** Amino acid sequences of the AncZB (blue) and AncBcd (orange) HDs (Liu et al, 2018). The four subdomains are labeled above the corresponding residues. Missing residues in the AncBcd sequence indicate identity with AncZB. Diagnostic residues conserved in all known Bcd HDs are labeled with asterisks. **B.** Schematic of the experimental design. Chimeric HDs between AncZB and AncBcd were inserted into the coding sequence (cds) of a Bcd rescue transgene. Shown below are the results of three preliminary experiments from Liu et al (2018), which show that the AncZB has no rescue activity (left), the AncZB HD with a double substitution (K50R54) provides partial rescue (middle), and the AncBcd HD provides full rescue activity. Transcripts whose expression patterns are represented are *hunchback* (*hb*), *giant* (*gt*), *orthodenticle* (*otd*), and *even-skipped* (*eve*). Blue circles represent abdominal segments (A1-A8), green circles represent thoracic segments (T1-T3), and red, brown and yellow circles represent head segments. Segments that give rise to wings and legs in the adult are shown. Filzkörper (Fk) are posterior larval structures.

In this paper, we present experiments designed to identify these other substitutions. We show that other substitutions in the RH collectively and synergistically contribute to HD function by increasing the number of target genes regulated by the AncZB HD. However, RH substitutions alone cannot fully rescue embryos lacking Bcd to adulthood. High frequency survival to adulthood is observed only if forward substitutions in the RH are combined with substitutions in two other subdomains (NT and H1). In contrast, combining substitutions in the RH with those in helix H1 or NT alone generates suboptimal HDs that achieve full rescue to adulthood, but with much lower frequencies. Taken together, these results suggest a multi-step pathway to explain the evolutionary transition from a non-functional AncZB HD to an AncBcd HD with robust *in vivo* function.

## Results

### Epistasis between amino acids in the RH increased the activity of the AncBcd HD

Among the 31 amino acid differences between the AncZB and AncBcd HDs, 11 are highly conserved in all available Bcd HD sequences, and are not found in any available Zen HD sequences [[34]; Figure 1A]. Six of these “diagnostic” substitutions, including q50>**K** and m54>**R**, are present in the RH subdomain, which directly contacts base pairs in the major groove of DNA [38]. We hypothesized that one or more diagnostic RH substitutions besides q50>K and m54>R might augment the degree of rescue mediated by the AncZB_**K50R54** HD. To start, we added all four (residues 42, 43, 55, and 58) to the AncZB_**K50R54** HD to generate the AncZB_**RHdiag** HD (Figure 2I-L). Surprisingly, when inserted into a *bcd* rescue transgene, the AncZB_**RHdiag** HD showed a lower level of rescue than the AncZB_**K50R54** HD. For example, no head structures were observed in larvae carrying the AncZB_**RHdiag** HD, and only 40% formed two thoracic segments [compared to 80% for the AncZB_**K50R54** (Figure 2I, J, compared to Figure 2E, F)]. This lower level of rescue activity was also observed at the transcriptional level. Consistent with the missing head structures, expression of the head gap gene *otd* [easily detectable in embryos rescued by the AncZB_**K50R54** construct (Figure 2H)] was not detected in embryos rescued with the AncZB_**RHdiag** construct (Figure 2L). We also observed reductions in the expression patterns of *hb* and *gt*, and anterior shifts of these patterns compared to those activated by the AncZB_**K50R54** HD (Figure 2L, compared to 2H). For *hb*, we quantified this shift by measuring the posterior boundary position (pbp) as a percentage of embryo length (% EL, where 100% = the anterior tip) (see Methods). In embryos carrying the AncZB_**RHdiag** embryos, the average position was at 82% EL (Figure S1B), while the average PBP for the AncZB_**K50R54** construct was at 77% EL (Figure S1A). Finally, the AncZB_**RHdiag** construct did not detectably repress Cad translation (Figure 2K).

**Figure 2:**
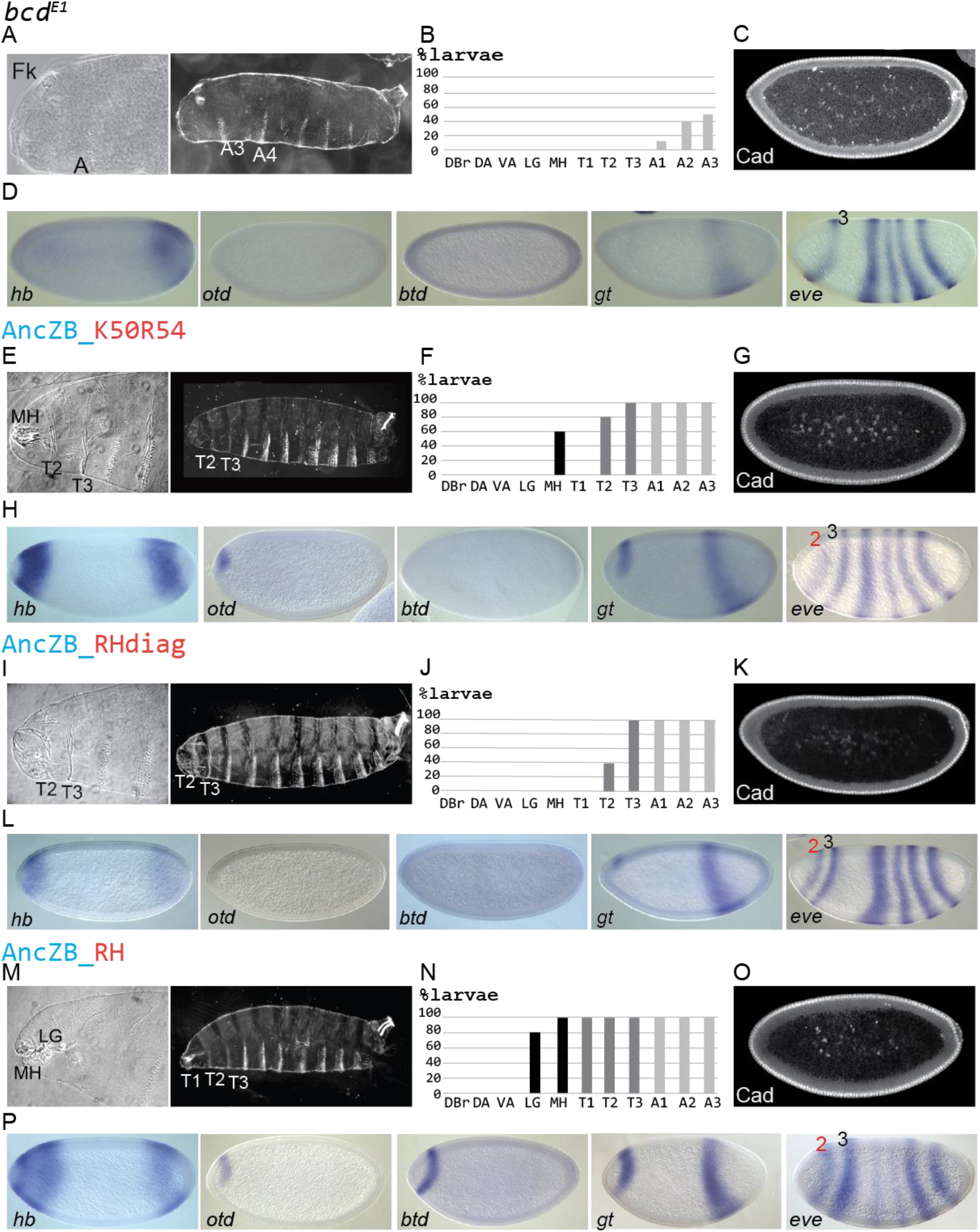
Morphological and molecular activities provided by diagnostic and nondiagnostic changes in the RH. Morphological structures and molecular activities are shown for embryos lacking Bcd (*bcd*^*E1*^; **A-D**), and in embryos rescued with by AncZB_**K50R54** (**E-H**), AncZB_**RHdiag** (*I-L*), and AncZB_**RH** (**M-P**). For each experiment, cuticle preparations of first instar anterior regions and whole larvae are shown (**A, E, I, M**), along with the percentages of first instar larvae that formed specific morphological structures (**B, F, J, N**). Indicated structures include Filzkoerper (Fk), Dorsal bridge (DBr), Dorsal Arm (DA), ventral arm (VA), lateralgraete (LG), mouth hooks (MH), the three thoracic segments (T1-T3), and three anterior-most abdominal segments (A1-A3). **C, G, K, O.** Caudal (Cad) immunostaining in representative stage 5 embryos. **D, H, L, P.** Patterns of Bcd target genes *hunchback* (*hb*), *orthodenticle* (*otd*), *buttonhead* (*btd*), *giant* (*gt*), and *even-skipped* (*eve*) in representative stage 5 embryos. Measurements of anterior *hb* patterns are shown in Figure S1.

These experiments suggest that negative epistatic interactions exist among diagnostic residues in the AncZB_**RHdiag** HD, which reduce biological activity compared to the AncZB_**K50R54** double substitution. One possibility is that these negative interactions are mitigated by the other three non-diagnostic substitutions in the AncBcd RH (Figure 1). To test this, we replaced the whole RH from AncZB HD with that of AncBcd (AncZB_**RH**) (Figure 2M-P). The addition of three more substitutions in the RH substantially improved the *in vivo* activity compared to both the AncZB_**K50R54** and the AncZB_**RHdiag** constructs. Around 95% of larvae containing AncZB_**RH** formed all three thoracic segments, and more than 80% formed cephalic structures (mouthhooks (MH) and lateralgraete (LG) only; Figure 2M, N). However, no larvae carrying the AncZB_**RH** construct survived to adulthood. At the molecular level, early embryos activated transcription of the target genes *otd* and *btd* (Figure 2P), which were not activated by the AncZB_**RHdiag** (Figure 2L), but failed to activate *eve* stripe 1 (Figure 2P). Also, the expression patterns of *hb* and *gt* were more strongly activated in AncZB_**RH** embryos compared to AncZB_**RHdiag**. In particular, the average *hb* pbp in AncZB_**RH** embryos was at 73% EL (Figure S1C), which is more posteriorly localized compared to 82% EL in AncZB_**RHdiag** embryos (Figure S1B). We could not detect any significant repression of Cad translation in AncZB_**RH** embryos (Figure 2K).

Taken together, these results suggest that positive and negative epistatic interactions within the RH were critical for the evolution of the AncBcd HD. However, none of the RH substitutions tested here mediate full rescue of Bcd deficient embryos to adulthood, indicating that additional substitutions in other subdomains were required for the acquisition of Bcd’s novel patterning activities.

### Combining substitutions in three subdomains were required for the evolution of robust AncBcd HD function

We tested several different constructs that combine forward substitutions in the RH with those in other subdomains. In these experiments, the cuticle patterns of first instar larvae containing each construct were highly variable, so we divided them into the following four categories (see Methods): 1. WTL (wild type like): larvae with easily detectable head structures (MH, LG, VA, DA, and DBr), three thoracic segments, and eight abdominal segments. 2. ΔHead: larvae missing any of the five head structures mentioned above. 3. ΔHead + Abdomen: larvae with head defects and additional defects in abdominal segments. 4. ΔAbdomen: larvae with normal head structures, but with defects in abdominal segments. As positive controls, we assayed the rescue activities of transgenes containing the wild type *Drosophila* Bcd HD and the reconstructed AncBcd HD, which produced ~90% and ~65% WTL larvae, respectively (Figure 3A, B; S2A and B). For the wild-type transgene, the remaining 10% were classified as ΔHead, and no larvae showed abdominal defects. In contrast, larvae rescued with the AncBcd HD that were not classified as WTL showed more variability, with around 20% with head defects alone or a combination of head and abdominal defects. An additional 15% contained well-formed head structures, and severe defects in abdominal segments, which ranged from a mild phenotype showing only four segments to a strong phenotype lacking all abdominal segments and poorly formed filzkörper (Figure S2B).

**Figure 3:**
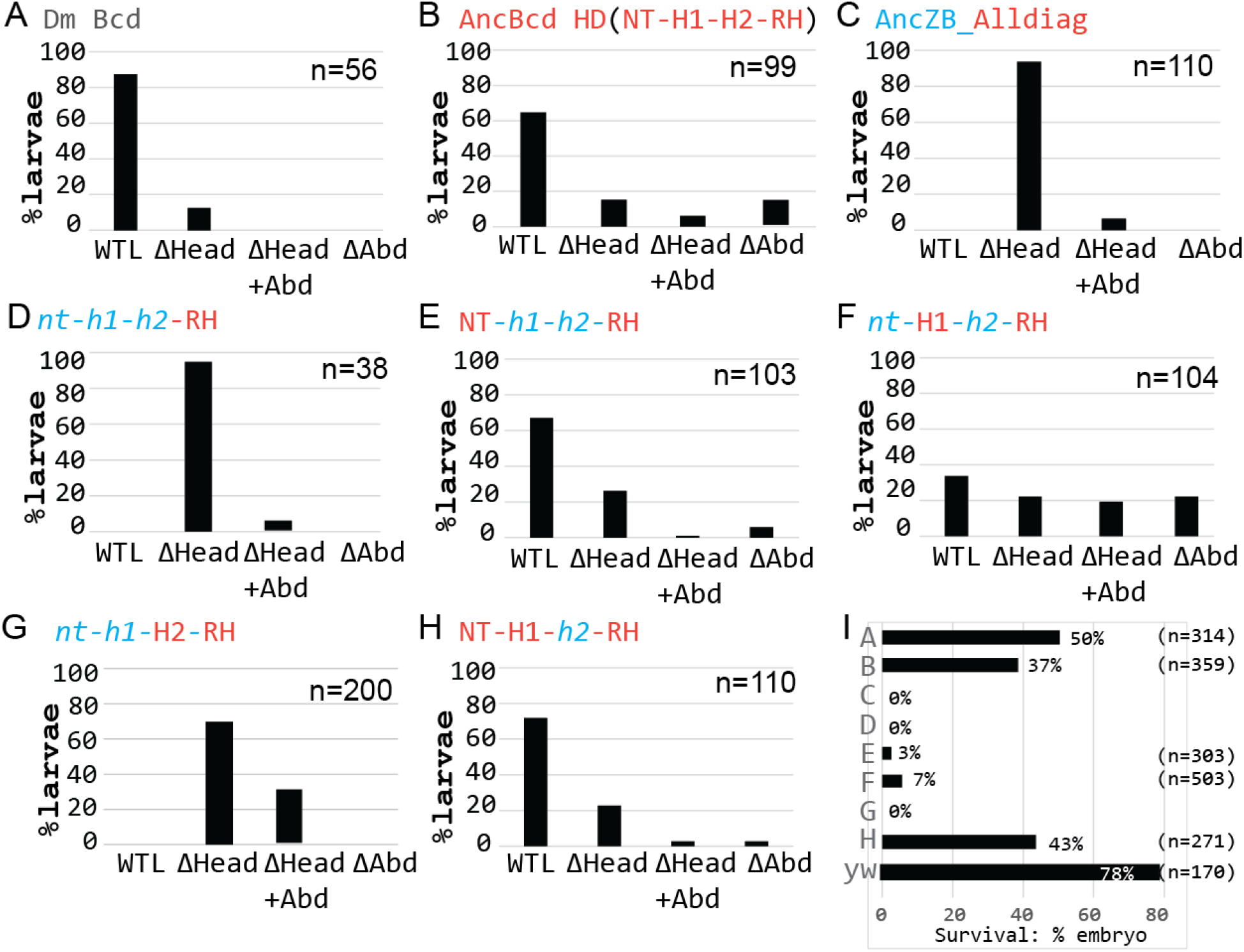
Different phenotypes observed among Drosophila larvae expressing ancestral HD proteins. Percentages of Bcd-deficient first instar larvae rescued with the indicated transgenes (**A-H**) that appear wild type-like (WTL), or have head defects (ΔHead), head plus abdominal defects (ΔHead+Abd), or abdominal defecvrts alone (ΔAbd). See Figure S2 for images of representative embryos in each phenotypic category. **I.** Survival rates of embryos to adulthood in the corresponding transgenic lines.

The ancestral reconstructions of the AncZB and AncBcd HDs identified 11 diagnostic changes distributed across all four subdomains (Figure 1A). We hypothesized that forward substitutions at all 11 diagnostic positions might convert the inactive AncZB HD into a fully active HD. Thus we made all 11 substitutions in the AncZB HD (AncZB_**Alldiag**), and tested it for rescue activity (Figure 3C). These substitutions in multiple subdomains showed partial patterning activity, with more than 90% of larvae forming all three thoracic segments, and more than 80% forming at least one of the head structures mentioned above. However, no larvae rescued by the AncZB_**Alldiag** transgene formed all five assayed head structures, so none could be classified as WTL, and no larvae survived to adulthood (Figure 3I).

We next tested the rescue activity of three chimeric HDs that separately combine all forward substitutions (diagnostic and non-diagnostic) in the RH with those in each of the other subdomains [**CH** (**NT**-*h1*-*h2*-**RH**)], [CH(*nt*-**H1**-*h2*-**RH**)] and [CH(*nt*-*h1*-**H2**-**RH**)]. Transgenes containing two of these chimeric HDs strongly increased rescue activity compared to the transgene containing all changes in the RH alone (Figure 3E, F compared to Figure 3D). Nearly 70% of *bcd* mutant larvae carrying the **CH** (**NT**-*h1*-*h2*-**RH**) transgene were classified as WTL (Figure 3E), but in many cases, specific head structures, including the lateralgraete and the dorsal and ventral arms, appeared shorter than normal (Figure S2E). An additional 25% failed to form one or more head structures. For the CH(*nt*-**H1**-*h2*-**RH**) transgene, there was also a strong increase in rescue activity, but less than 40% were classified as WTL, with the rest evenly distributed among the other three phenotypic categories (Figure 3F, and S2F). In contrast, no WTL larvae were produced by the CH(*nt*-*h1*-**H2**-**RH**) rescue transgene (Figure 3G, and S2G).

We performed hatching tests (see Methods) to monitor the frequency of survival past larval stages for the experiments that produced WTL larvae (Figure 3I). As a baseline, the frequency of survival to adulthood for wild type larvae under our laboratory conditions was ~80% (Figure 3I). In contrast, the positive control transgenes containing endogenous Bcd and AncBcd HDs resulted in the survival of only 50% and 37% of larvae to adulthood respectively, perhaps due to the fact that the transgenes are inserted into an ectopic genomic position. Remarkably, both chimeric constructs that yielded WTL larvae [CH(**NT**-*h1-h2*-**RH**) and CH(*nt*-**H1**-*h2*-**RH**)] directed the survival of 3% and 7% of those larvae to adulthood, respectively (Figure 3I). While these survival frequencies are quite low compared to the control experiments, they show that the full developmental function of the Bcd HD can be achieved by substitutions in two different combinations of subdomains (NT+RH and H1+RH).

We also tested if combining substitutions in the NT, H1, and RH subdomains would increase the rate of survival to adulthood. As predicted, 60% of larvae produced by *bcd* females containing the CH(**NT**-**H1**-*h2*-**RH**) were classified as WTL (Figure 3H; Figure S2H), and 43% survived to adulthood (Figure 3I), results that are very similar to those obtained by the AncBcd HD. This result shows that combining substitutions in three separate subdomains is required and sufficient for generating a Bcd HD with high penetrance rescue activity.

### Bcd target gene positions that correlate with full rescue to adulthood

To begin to understand the molecular basis for the differential rescue mediated by the chimeric HDs, we examined the expression patterns of several Bcd target genes, starting with Cad, which is translationally suppressed by Bcd in wild type embryos. As expected, all three chimeric HDs that direct full rescue also show Cad suppression (Figure S3C-E). In contrast, strong suppression of Cad was not detected in embryos partially rescued by the CH(*nt-h1*-**H2-RH**) transgene. However, our experiments with HDs containing RH substitutions alone or all diagnostic substitutions showed that one of these partially rescuing constructs (AncZB_**AllDiag**) also suppressed Cad (Figure S3A). Taken together, these results show that suppression of Cad may be required for rescuing to adulthood, but it is not sufficient.

We next examined *hb* expression in embryos containing the chimeric HD transgenes (Figure 4A, D, G, J, M, P). Our experiments with HDs containing RH substitutions alone or all diagnostic substitutions showed that the degree of partial rescue activity is positively correlated with the extension of the *hb* expression domain into middle regions of the embryo (Figure S1). However, the *hb* posterior boundary position (pbp) in the best case (AncZB_**Alldiag**) is located at 70%EL, relatively far from the boundary position in embryos rescued by the AncBcd HD (54% EL) (Figure S2D and E). Thus we hypothesized that HDs capable of directing full rescue to adulthood might activate *hb* domains that extend farther posteriorly than those that fail to rescue. Indeed, embryos containing all three fully rescuing constructs show *hb* pbps that range from 68 to 63% EL (Figure 4G, J, M). However, the correlation between *hb* boundary positioning and full rescue is not perfect. Specifically, the CH(*nt-h1*-**H2-RH**) chimera, which completely failed to fully rescue (Figure 3I), activated a *hb* domain with a pbp at 67% EL (Figure 4D). Therefore, these results suggest that extending the *hb* domain to a specific AP position is also required, but not sufficient for the mediating the full regulatory activity of the AncBcd HD.

**Figure 4:**
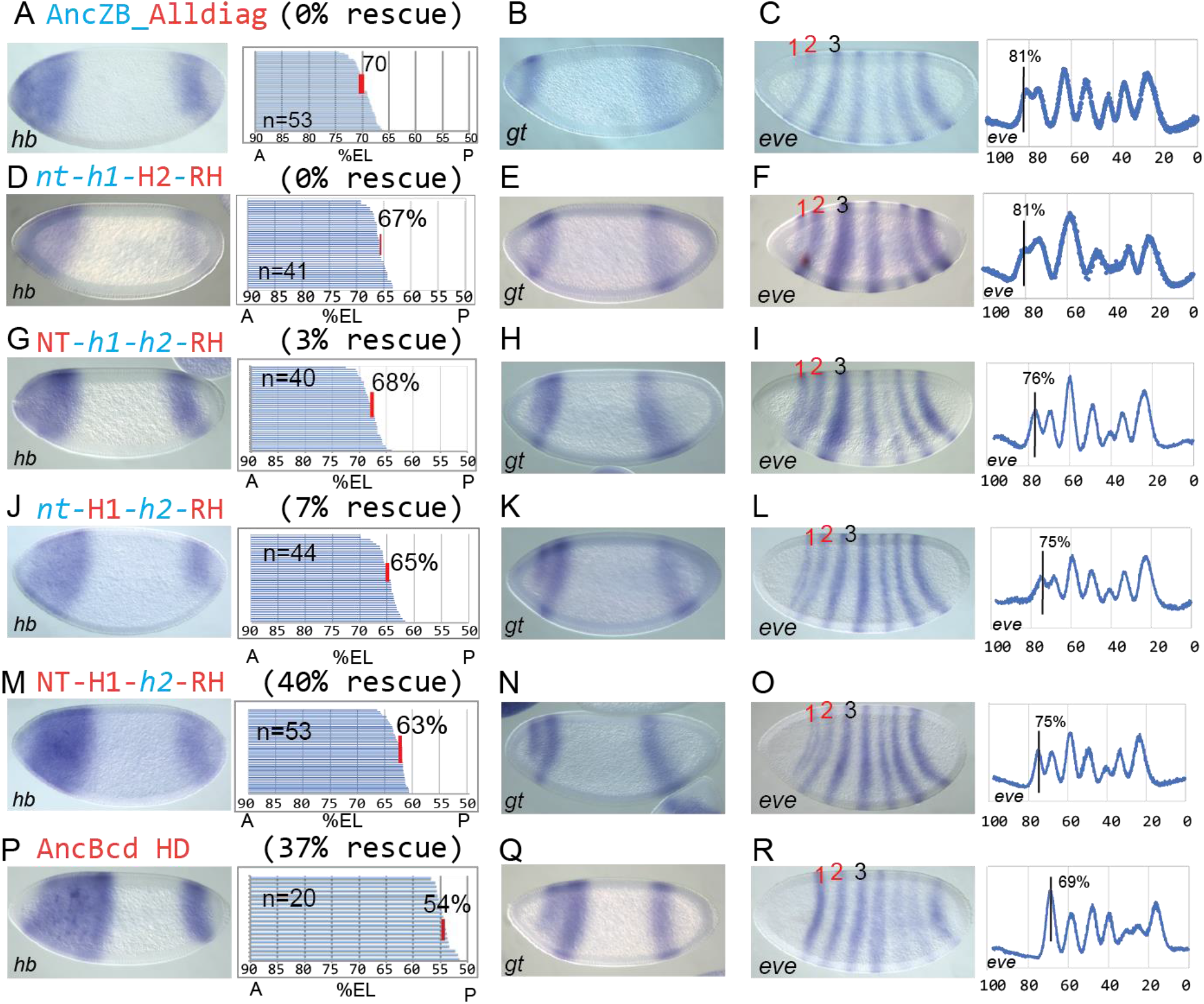
hb, gt and eve expression patterns in lines exhibiting different levels of phenotypic rescue. Tested HDs are labeled as in Figure 3. **A, D, G, J, M, P.** Representative stage 5 embryos stained by *in situ* hybridization to detect *hb*. Panels show the analysis of multiple embryos to calculate the average posterior boundary position (% EL; anterior tip = 100%) of *hb* at stage 5. Each horizontal line in each panel represents the anterior *hb* expression pattern in a single embryo, and the average pbp is denoted by a vertical red line. **B, E, H, K, N, Q.** Representative stage 5 embryos stained for *gt* expression. **C, F, I, L, O, R.** Representative stage 5 embryos stained for *gt* expression. *eve* stripe 1 positions were calculated from more than 10 individual embryos and are denoted by vertical black lines.

We also examined the expression of the gap genes *otd*, *btd*, and *gt*, and the pair-rule gene *eve* (Figure 4 and Figure S3). The expression patterns of *otd* and *btd* were indistinguishable in embryos carrying the three chimeric transgenes that fully rescue (Figure S3C-E) and the one that does not (Figure S3B). In contrast, the *gt* expression pattern showed significant differences. The anterior *gt* expression domain initially appears as a broad stripe, which resolves over time into two stripes [39]–[41]. The separation into stripes occurs in all three lines that direct full rescue (Figure 4H, K, N), but not in embryos carrying the CH(*nt-h1*-**H2**-**RH**) chimera (Figure 4E), or in any other tested constructs that fail to direct full rescue (Figure 2H, L, P and Figure 4B). We also observed a strong correlation between fully and partially rescuing chimeric lines and the positioning of the anterior-most *eve* stripes. Embryos carrying the three transgenes that fully rescue formed *eve* stripe 1 at 75-76% EL (Figure 4I, L, O), while the CH(*nt*-*h1*-**H2**-**RH**) and AncZB_**Alldiag** transgenes (both 0% full rescue) consistently formed this stripe more anteriorly (81% EL; Figure 4C and F). Also, *eve* 1 was more clearly separated from *eve* 2 in embryos that fully rescue to adulthood. Thus, there is a perfect correlation between the ability to fully rescue to adulthood, the separation of the anterior *gt* domain into two stripes, and the positioning of the anterior-most *eve* stripes. However, the positions of the *gt* domain and *eve* stripe 1 even in these fully rescuing lines were still significantly anterior compared to the control AncBcd HD line (Figure 4I, L and O compared 4R).

Although we observed substantial differences in *gt* and *eve* patterning between fully and partially rescuing lines, we detected only one slight expression difference that might explain the different survival rates (3-40%) among the three constructs that fully rescue. Notably, the **CH(NT-H1**-*h2*-**RH**) transgene, which combines forward substitutions in three subdomains and rescues 40% of *bcd* mutant embryos to adulthood activated *hb* expression with a pbp at 63% (Figure 4M). This position is slightly posterior compared to the boundaries in embryos rescued by the **CH** (**NT-***h1*-*h2*-**RH**) or **CH** (*nt***-H1**-*h2*-**RH**) transgenes (68% and 65%, respectively, Figure 4G, J). Aside from this difference, we detected no changes in any of the tested gap gene or *eve* expression patterns among these constructs.

## Discussion

### Molecular requirements for the patterning activity of the AncBcd HD in Drosophila

In this paper, we used an *in vivo Drosophila* rescue assay to study the impact of the historical coding sequence changes on the evolution of Bcd HD’s developmental functions. By making chimeric HDs between the AncZB (no function) and the AncBcd (full function) HDs, we showed that the substitutions in at least three separate subdomains (NT, H1, and RH) must be combined for full patterning activity.

AncBcd evolved to suppress translation of Cad and activate transcription of a large number of target genes at different positions along the AP axis of the embryo. Our results shed light on the molecular requirements for both of these activities. The R54 residue in Bcd was previously shown to be required for Cad suppression [31], but our data suggest that it is not sufficient, even in combination with the other eight forward substitutions in the RH of AncBcd (AncZB_**RH**). However, by combining the RH substitutions with several different sets of substitution in the NT and/or H1 or substituting all diagnostic residues across all subdomains, variable levels of suppression were achieved (Figure S3). The impact of the level of suppression on the rescue potential and patterning is not known, and will be addressed by future experiments.

At the transcriptional level, it was previously shown that inserting K50 alone into the AncZB HD caused the activation of only three of eight tested target gene responses, while the double substitution (K50R54) increased that number to five [34]. In this paper, we show that substituting all nine amino acids from the RH of AncBcd HD into AncZB resulted in the activation of all tested target genes with missing patterns (*eve stripe* 1 and anterior stripes of *gt*). Moreover, these Bcd-dependent expression patterns were anteriorly shifted (Figure 2P). Combining substitutions in the RH with those in NT and H1 had major effects on the gene expression patterns: they extended or shifted critical expression patterns into more posterior positions, which might have allowed for splitting of the anterior *gt* domain into the two-striped pattern seen in wild type embryos.

The correlation between target gene expansion and rescue activity is most easily observed for the target gene *hb*, which encodes a critical cofactor for activation of all Bcd-dependent target genes [42]–[45], and functions as an important repressor to prevent posterior gap gene expression in anterior regions of the embryo [46]–[49]. In embryos carrying constructs that fail to fully rescue to adulthood, *hb* pbps are located between 82 to 67% EL, while embryos carrying constructs with full rescue activity form *hb* pbps at the posterior limit of this range (68% EL) or farther posterior. Interestingly, the [**CH** (**NT-H1**-*h2*-**RH)]** construct, which rescues to adulthood with a frequency similar to that observed for the AncBcd HD control, forms a *hb* pbp at 63%.

We propose that the position of ~65% EL establishes the minimal amount of embryonic space required for establishing the correct placement of gap and pair-rule stripes, robust formation of cephalic structures, and ultimately survival to adulthood. The pbp at 65% EL is significantly more anterior than those directed by the control AncBcd construct (54% EL, Figure 4K) or wild type embryos (54%; [29], but is very close to the *hb* pbp in embryos laid by heterozygous *bcd* females (~61% EL), which survive with high penetrance [50]. How interactions between the RH and other HD subdomains cause posterior extensions of the zygotic *hb* domain is not clear; they could indirectly modify the DNA-binding preferences of the HD or mediate interactions with maternal cofactors such as Hb or Zelda, both of which are critical for Bcd’s *in vivo* functions in *Drosophila* [42], [43], [51]–[54].

### Evolution of the AncBcd HD through suboptimal intermediate steps

An ancient duplication of AncZB led to the evolution of the K50 HD protein Bcd as a key regulator of anterior development in the Cyclorrhaphan suborder of the Diptera (two-winged insects) [21]. None of the other suborders of the Diptera or other insects contain Bcd; in these insects, maternal Bcd’s roles in anterior patterning must be fulfilled by other gene(s). In the flour beetle *Tribolium* and the jewel wasp *Nasonia*, Bcd-like activity is partially provided by maternal Orthodenticle (Otd), another K50 HD transcription factor that binds to DNA sequences similar to those bound by Bcd [55], [56]. Because many Bcd target genes, unlike Bcd, are highly conserved, it has been proposed that Bcd evolved to take over regulation of an ancestral network of genes regulated by a protein with similar specificities, such as K50 protein Otd [55]. In *Drosophila*, *otd* has evolved to become a Bcd target gene [57], and its ancestral role in anterior patterning has been diminished [58]. In this evolutionary scenario, Bcd might have gained anterior patterning role through stepwise changes in the coding sequence, which led to increased DNA-binding activities and modified DNA-specificities. However, there are other ways and unrelated proteins [a homolog of Odd-paired in the drain fly *Clogmia*, and a cysteine clamp protein in the midge *Chironomus*] as the maternal anterior patterning factor [59], [60], suggesting alternative ways of anterior segmentation among flies and insects.

Our data show that robust patterning function of the AncBcd from AncZB is achieved by combining forward substitutions in three subdomains (RH, NT, and H1). It seems impossible that critical amino acid substitutions in all three subdomains occurred simultaneously at some point in the evolution of the AncBcd HD. However, our data suggest that critical substitutions in each of the three might have occurred in a specific temporal order, each of which endowed the protein with a novel property that could be positively selected in evolving flies (Figure 5).

**Figure 5:**
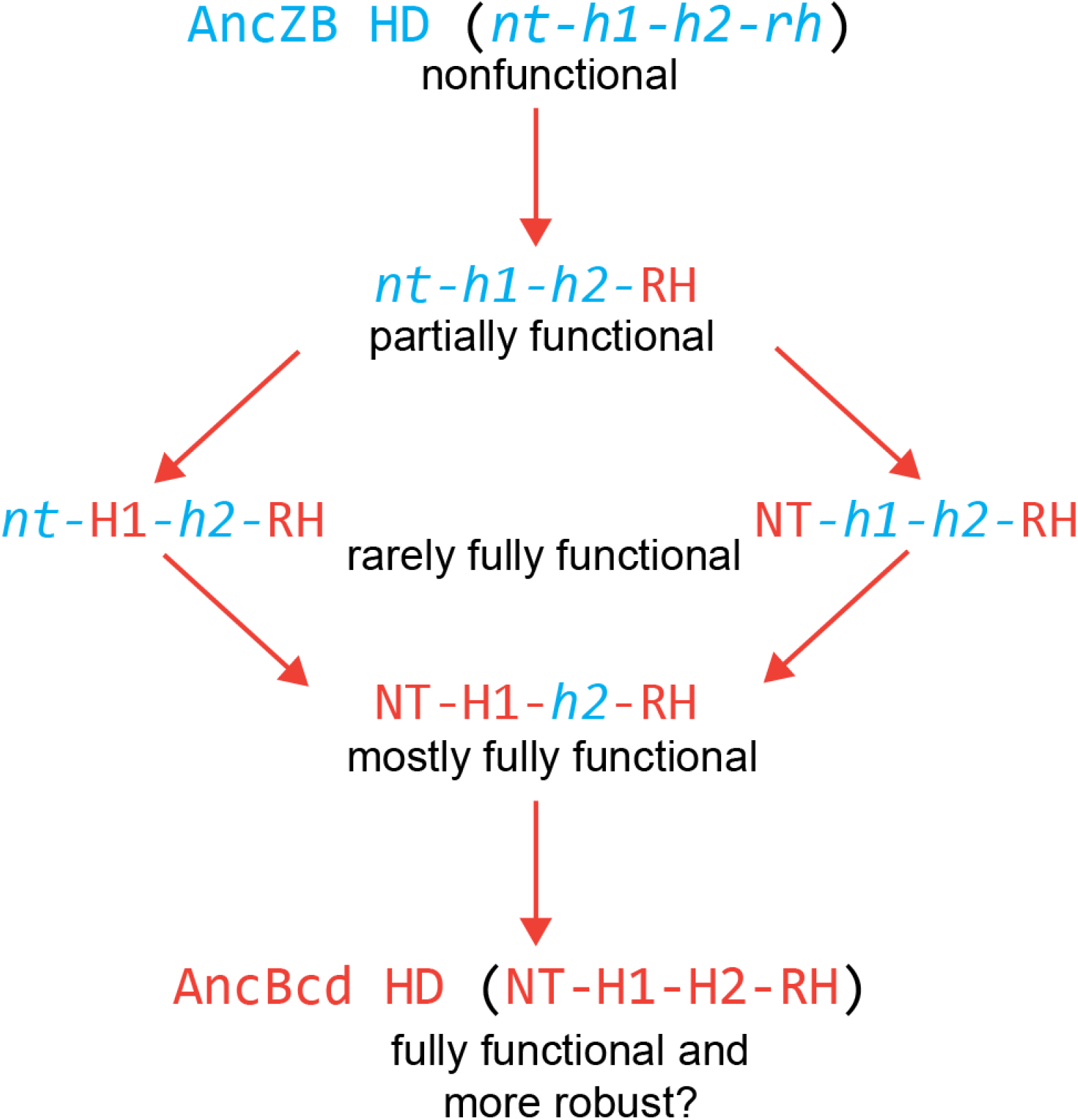
A proposed multi-step pathway for the evolution of the AncBcd HD. Orange arrows represent amino acid substitutions in individual subdomains. In the first step, initial substitutions in the RH changed the DNA-binding preferences of the HD, and allowed it to bind to RNA. In step 2, these initial substitutions were followed by additional changes in either the NT or the H1 subdomain, each of which could have significantly augmented the *in vivo* activities of the evolving HD in a small percentage of embryos. In a third step, substitutions in the unchanged subdomain (H1 for RH+NT or NT for RH+H1) would further increase patterning activity and raise the survival rate to almost control levels.

Assuming that the ancestral network was controlled by a K50 HD protein such as Otd [58], we propose that the first step involved multiple substitutions in the RH, including q50>K and m54>R. The codons for Q (Gln: CAA and CAG) and K (Lys: AAA and AAG) differ by only one base, so the q50>K transition involved only a single base pair substitution that would have dramatically changed the evolving protein’s DNA-binding preference. Reverse substituting or mutating K50 completely abolishes AncBcd HD function[34], which means that the effects of all other substitutions in the evolving HD were dependent on keeping the K50 residue intact. If this substitution occurred in an ancestral fly with an Otd-dependent anterior patterning network, the evolving protein would be immediately available to bind to many Otd-dependent target genes, which might have provided a selective advantage. The m54>R substitution, which also involves a single base change (AUG to AGG), might have refined DNA-binding specificity to increase the number of activated target genes, and set the stage for other substitutions that allowed the AncBcd HD to bind to RNA. K50 and R54 are present together only in Bcd HDs [37], consistent with the possibility that this combination might have been under positive selection. In addition to the q50>K and m54>R substitutions, there are seven other amino acid differences between the RH subdomains of AncZB and AncBcd. It is not clear which of these are required for AncBcd HD function, or when they appeared historically. However, one combination of six substitutions tested here (AncZB_**RHdiag**) reduced HD activity compared to the K50R54 double substitution. This result suggests that interactions between amino acids constrained the historical order of substitutions in the RH subdomain.

While robust HD activity requires substitutions in three subdomains, forward substitutions in either NT or H1 substantially augment the rescue activity generated by changes in the RH alone. Specifically, AncZB HDs containing either combination (RH+NT or RH+H1) rescue a small percentage of embryos that survive to adulthood (Figure 3I). We propose that the addition of substitutions in either NT or H1 represent alternative second steps in the historical evolution of AncZB HD (Figure 5). Either combination (RH +NT or RH+H1) would have generated a suboptimal intermediate HD configuration that could have been positively selected for and stabilized, perhaps by increasing the fitness of a sub-population in specific physical/environmental conditions. Once stabilized, in a third step, substitutions in the other critical subdomain (H1 for the NT+RH intermediate, for example) would further increase HD activity and robustness of the evolving HD.

### Limitations and challenges for the future

Our results shed light on the mechanisms involved in the evolution of the AncBcd HD, but are limited by the fact that all chimeric HDs were inserted into the modern-day *Drosophila* Bcd protein. As such, these experiments do not take into account the evolution of other parts of the protein, which show even greater levels of amino acid sequence divergence. Further, all our experiments were performed in modern-day *Drosophila* embryos, and do not take into account changes in the cis-regulatory elements of target genes that co-evolved with the AncBcd protein. However, as the genome sequences of more insects become available, it should be possible to use reconstruction strategies to define the ancestral sequences of the complete AncBcd protein and the regulatory regions it interacts with. Furthermore, the ever-increasing use of CRISPR/Cas9 techniques for gene editing in non-model organisms should allow for testing ancestral protein and regulatory sequences in multiple insect species. While these methods cannot create the ancestral systems themselves, they should make it possible to discover general features that permit a transcription factor and its target regulatory sequences to co-evolve.

## Acknowledgements

We thank Danyang Yu for technical assistance, Rhea Datta and Patrick Lemaire for stimulating discussions, and Claude Desplan, Esteban Mazzoni, Erik Clark, and Matt Rockman for suggestions that greatly improved the manuscript. This project was supported by NIH Grant RO1 GM 51946 to SS.

## Author contributions

Conceptualization, P.O. and S.S.; Methodology, P.O., M.Z., J.L., and S.S.; Investigation, P.O., H.I.G., K.Y.U., and M.Z., T.T.; Writing – Original Draft, P.O. and S.S.; Writing – Review & Editing, P.O. and S.S..; Funding Acquisition, S.S.; Resources, M.Z. and J.L..; Supervision, P.O. and S.S.

## Methods

### *Drosophila* stocks, cloning, and transgenesis

We used the following stocks from our own lab for these experiments: yw (wild type), *Cyo bcd*^+^/*Sco*; *bcd*^*E1*^/*bcd*^*E1*^, yw; TM3B, Sb, Ser/D and ΦC31 (y+); 38F1 (w+). We cloned an injection plasmid (piattB40-Bcd) containing two inverted ΦC31-specific recombination sequences, a Gmr-GFP reporter and a polylinker flanked by 1.9kb *bcd* promoter and 0.8kb 3’UTR. The *bcd* coding region was amplified by PCR from pBS-SK+ cDNA clones, digested with RsrII and AscI and ligated into piattB40-Bcd in between Bcd promoter and 3’UTR. This main plasmid was used to generate Dm Bcd protein with different ancestral HDs, which are predicted as described and published in [34]. We used standard cloning techniques to generate homeodomain swaps and residue changes. Gene Blocks coding for the ancestral and chimeric HD sequences together with the flanking Bcd coding sequence were obtained from Integrated DNA Technologies (IDT). They were digested with AscI and BspEI and ligated to the piattB40-Bcd vector digested with the same restriction enzymes. The cloned sequences were confirmed by sanger sequencing before and after transfection. All transgenic lines were generated using the ΦC31 integration system (Recombination mediated cassette exchange, RMCE), and constructs were integrated into the 38F1 landing site on the second chromosome [61]. Each transgene was crossed to *Cyo bcd*^+^/*Sco*; *bcd*^*E1*^/*bcd*^*E1*^ to generate *Cyo bcd*^+^/*[transgene]*; *bcd*^*E1*^/*bcd*^*E1*^ stocks. Embryos and larvae from homozygous transgenic females were assayed for gene expression and cuticle phenotype.

### In situ hybridization, immunohistochemistry, and image processing

*In situ* hybridizations were performed as previously described [62]. Briefly, embryos 1-3 hours AEL (after egg laying) were dechorionated 2 minutes in 100% bleach, fixed and devitellinized in a biphasic fixation solution containing 3 ml 1X PBS, 1 ml 37% Formaldehyde and 4 ml Heptane for 25 minutes on a shaker at RT. Fixed and permeabilized embryos were incubated with DIG or Fluorescein-labeled RNA probes and the labeled probes were detected by Alkaline Phosphatase (AP)-conjugated primary antibodies (Roche Cat# 11093274910, RRID:AB_514497) and Roche Cat# 11426338910, RRID:AB_514504) by using NBT/BCIP solution (Roche Cat# 19315121). RNA expression was observed by Zeiss Axioskop microscopy.

Guinea pig anti-Cad [63] (1:400) and Alexa Fluor conjugated 647 donkey anti-guinea pig (1:500) (Molecular Probes Cat# A-21447, RRID:AB_141844) were used to examine Cad protein expression. All antibodies were diluted in PBT (1X PBS with 0.1% Tween). Data for immunostaining images were collected on a Leica TCS SP8 confocal microscope using the Leica confocal analysis software.

### Larval Phenotype Analyses and Hatching Assays

Cuticle preparations were performed on embryos aged 24-30 hours at 25°C as previously described [64]. Briefly, larvae were dechorionated for 2 minutes in 100% bleach, and a 1:1 mixture of methanol and heptane was used to remove the vitelline membrane and fix the larvae. Then these larvae were mounted in 1:1 mixture of Hoyer’s medium [65] and lactic acid and incubated o/n at 65°C to digest inner tissues.

Dark-field views of whole larvae were imaged at 200X magnification; DIC images of cephalic regions are imaged at 400X. Each image was sorted into one of three categories that encompassed the variation in phenotype of first instar larvae both within each transgenic fly line, and across all fly lines analyzed. The different phenotypic categories are; WTL: larvae containing all head segments [MH, LG, VA, DA, and DBr (Dorsal Bridge), three thoracic segments, and eight abdominal segments], ΔHead: larvae missing one or more head segments with normal abdominal and thoracic segments, ΔAbdomen: larvae with variable defects in abdominal segments but normal head and thorax. If an embryo showed both abdominal defects and head defects, it was classified as ΔHead+Abd. The number of WTL, ΔHead, ΔHead+Abd and ΔAbdomen larvae were counted for each transgenic line, tabulated and graphed as a percentage of the total number of embryos analyzed for that transgenic line.

With the lines that we observed WTL larvae, we set up hatching assays to assess survival of these larvae to pupa stage and then adulthood. For hatching assays flies were let lay eggs on fruit juice plates for an hour and then over 100 eggs were picked and incubated until the pupae are formed and pupae were counted as the survival rate per the embryos picked.

### Measuring hb Posterior Boundary Positions (PBPs)

To measure PBPs of *hb* anterior expression, stained embryos of appropriate ages were imaged at 200× on a Zeiss Axioskop. Briefly, coordinates were established for each embryo so that the x- and y-axes were tangential to the ventral and anterior sides, respectively. A-P positions were displayed as percent of embryo length (EL%) with the anterior pole as 100%. PBPs were determined by visual estimation as the distance from the anterior tip to the most posterior position of *hb* anterior expression and these results were confirmed by ImageJ (ImageJ, RRID: SCR_003070) [66], [67] analyses.

Embryo images were loaded into ImageJ, and a Region of Interest (ROI) that was approximately 35% width of DV length from 95% to 60% (where 100% is most dorsal side) to analyze expression patterns was generated. The width of the ROI was kept constant when imaging all embryos, but the length was varied such that the length of the ROI spanned the length of the whole embryo. For each embryo, an intensity profile plot (intensity v. position along embryo length) was generated for the ROI. The midpoint of the curve that represents the edge of the boundary of expression of target gene was selected as the position of the boundary of gene expression. This numerical position was divided by the total embryo length to normalize the PBPs by % EL, and PBPs from individual embryos were averaged. 100% EL denotes the anterior tip, and 0% represents the posterior tip of the embryo.

### Measuring eve patterns

To measure *eve* stripe patterns, stained embryos at stage 5 were positioned as described above and imaged at 200X. For each embryo, an intensity profile plot (intensity vs position along embryo length, where 100% denotes the anterior tip) was generated using ImageJ (ImageJ, RRID: SCR_003070) [66], [67], and analyzed using our Embryo Analyzer tool, which serves to take ImageJ plots of fly embryos, and convert them to produce a single file of normalized intensities along the AP axis of n number embryos.

### Resource Availability

#### Lead contact

Stephen Small

#### Materials availability

All unique materials generated in this study are available upon request.

#### Data and code availability

The code for *hb* and *eve* stripe pattern measurement supporting the current study have not been deposited in a public repository because of the specificity of the measurement to our case but are available from the corresponding author on request.

Further information and requests for resources and reagents should be directed to and will be fulfilled by the Lead Contact, Dr. Stephen Small

## Notes

### Competing Interest Statement

The authors have declared no competing interest.

### Summary of Updates

Manuscript revisited figures updated to clarify our point. Conclusions remained the same.

## References

[1] I. S. Peter and E. H. Davidson, Genomic Control Process: Development and Evolution. 2015.

[2] B. Prud’homme, N. Gompel, and S. B. Carroll, “Emerging principles of regulatory evolution,” Proc. Natl. Acad. Sci. U. S. A., vol. 104, no. SUPPL. 1, pp. 8605–8612, May 2007, doi: 10.1073/pnas.0700488104.

[3] G. A. Wray, “The evolutionary significance of cis-regulatory mutations,” Nat. Rev. Genet., vol. 8, no. 3, pp. 206–216, Mar. 2007, doi: 10.1038/nrg2063.

[4] I. S. Peter and E. H. Davidson, “Evolution of gene regulatory networks controlling body plan development,” Cell, vol. 144, no. 6. Cell, pp. 970–985, Mar. 18, 2011, doi: 10.1016/j.cell.2011.02.017.

[5] G. P. Wagner and V. J. Lynch, “The gene regulatory logic of transcription factor evolution,” Trends in Ecology and Evolution, vol. 23, no. 7. Trends Ecol Evol, pp. 377–385, Jul. 2008, doi: 10.1016/j.tree.2008.03.006.

[6] C. Sayou et al., “A promiscuous intermediate underlies the evolution of LEAFY DNA binding specificity.,” Science, vol. 343, no. 6171, pp. 645–8, Feb. 2014, doi: 10.1126/science.1248229.

[7] S. Nakagawa, S. S. Gisselbrecht, J. M. Rogers, D. L. Hartl, and M. L. Bulyk, “DNA-binding specificity changes in the evolution of forkhead transcription factors,” Proc. Natl. Acad. Sci. U. S. A., vol. 110, no. 30, pp. 12349–12354, Jul. 2013, doi: 10.1073/pnas.1310430110.

[8] S. Ohno, Evolution by Gene Duplication. Berlin, Heidelberg: Springer Berlin Heidelberg, 1970.

[9] F. A. Kondrashov, I. B. Rogozin, Y. I. Wolf, and E. V Koonin, “Selection in the evolution of gene duplications,” Genome Biol., vol. 3, no. 2, p. research0008.1, Jan. 2002, doi: 10.1186/gb-2002-3-2-research0008.

[10] L. N. Singh and S. Hannenhalli, “Functional diversification of paralogous transcription factors via divergence in DNA binding site motif and in expression,” PLoS One, vol. 3, no. 6, Jun. 2008, doi: 10.1371/journal.pone.0002345.

[11] R. O. Emerson and J. H. Thomas, “Adaptive Evolution in Zinc Finger Transcription Factors,” PLoS Genet., vol. 5, no. 1, p. e1000325, Jan. 2009, doi: 10.1371/journal.pgen.1000325.

[12] D. Vlad et al., “Leaf shape evolution through duplication, regulatory diversification and loss of a homeobox gene,” Science (80-.)., vol. 343, no. 6172, pp. 780–783, Feb. 2014, doi: 10.1126/science.1248384.

[13] F. R. Schubert, K. Nieselt-Struwe, and P. Gruss, “The Antennapedia-type homeobox genes have evolved from three precursors separated early in metazoan evolution,” Proc. Natl. Acad. Sci., vol. 90, no. 1, pp. 143–147, Jan. 1993, doi: 10.1073/PNAS.90.1.143.

[14] J. Zhang and M. Nei, “Evolution of Antennapedia-Class Homeobox Genes,” Genetics, vol. 142, no. 1, p. 295, 1996, Accessed: May 21, 2020. [Online]. Available: https://www.ncbi.nlm.nih.gov/pmc/articles/PMC1206958/.

[15] C. Kappen, K. Schughart, and F. H. Ruddle, “Two steps in the evolution of Antennapedia-class vertebrate homeobox genes,” Proc. Natl. Acad. Sci., vol. 86, no. 14, pp. 5459–5463, Jul. 1989, doi: 10.1073/PNAS.86.14.5459.

[16] D. Duboule and P. Dollé, “The structural and functional organization of the murine HOX gene family resembles that of Drosophila homeotic genes.,” EMBO J., vol. 8, no. 5, pp. 1497–1505, May 1989, doi: 10.1002/j.1460-2075.1989.tb03534.x.

[17] J. M. Greer, J. Puetz, K. R. Thomas, and M. R. Capecchi, “Maintenance of functional equivalence during paralogous Hox gene evolution,” Nature, vol. 403, no. 6770, pp. 661–665, Feb. 2000, doi: 10.1038/35001077.

[18] R. Galant and S. B. Carroll, “Evolution of a transcriptional repression domain in an insect Hox protein,” Nature, vol. 415, no. 6874, pp. 910–913, Feb. 2002, doi: 10.1038/nature717.

[19] M. Ronshaugen, N. McGinnis, and W. McGinnis, “Hox protein mutation and macroevolution of the insect body plan,” Nature, vol. 415, no. 6874, pp. 914–917, Feb. 2002, doi: 10.1038/nature716.

[20] F. Falciani et al., “Class 3 Hox genes in insects and the origin of zen.,” Proc. Natl. Acad. Sci., vol. 93, no. 16, pp. 8479–8484, Aug. 1996, doi: 10.1073/pnas.93.16.8479.

[21] M. Stauber et al., “The anterior determinant bicoid of Drosophila is a derived Hox class 3 gene.,” Proc. Natl. Acad. Sci. U. S. A., vol. 96, no. 7, pp. 3786–9, Mar. 1999, doi: 10.1073/pnas.96.7.3786.

[22] U. Schmidt-Ott, A. M. Rafiqi, and S. Lemke, “Hox3/zen and the Evolution of Extraembryonic Epithelia in Insects,” Springer, New York, NY, 2010, pp. 133–144.

[23] M. Stauber, A. Prell, and U. Schmidt-Ott, “A single Hox3 gene with composite bicoid and zerknullt expression characteristics in non-Cyclorrhaphan flies.,” Proc. Natl. Acad. Sci. U. S. A., vol. 99, no. 1, pp. 274–9, Jan. 2002, doi: 10.1073/pnas.012292899.

[24] S.-K. Chan and G. Struhl, “Sequence-specific RNA binding by Bicoid,” Nature, vol. 388, no. 6643, pp. 634–634, Aug. 1997, doi: 10.1038/41692.

[25] R. Rivera-Pomar, D. Niessing, U. Schmidt-Ott, W. J. Gehring, and H. Jacklë, “RNA binding and translational suppression by bicoid,” Nature, vol. 379, no. 6567, pp. 746–749, Feb. 1996, doi: 10.1038/379746a0.

[26] W. Driever and C. Nüsslein-Volhard, “A gradient of bicoid protein in Drosophila embryos.,” Cell, vol. 54, no. 1, pp. 83–93, Jul. 1988, [Online]. Available: http://www.ncbi.nlm.nih.gov/pubmed/3383244.

[27] W. Driever, G. Thoma, and C. Nüsslein-Volhard, “Determination of spatial domains of zygotic gene expression in the Drosophila embryo by the affinity of binding sites for the bicoid morphogen,” Nature, vol. 340, no. 6232, pp. 363–367, Aug. 1989, doi: 10.1038/340363a0.

[28] G. Struhl, K. Struhl, and P. M. Macdonald, “The gradient morphogen bicoid is a concentration-dependent transcriptional activator,” Cell, vol. 57, no. 7, pp. 1259–1273, Jun. 1989, doi: 10.1016/0092-8674(89)90062-7.

[29] H. Chen, Z. Xu, C. Mei, D. Yu, and S. Small, “A system of repressor gradients spatially organizes the boundaries of Bicoid-dependent target genes.,” Cell, vol. 149, no. 3, pp. 618–29, Apr. 2012, doi: 10.1016/j.cell.2012.03.018.

[30] A. Nasiadka, B. H. Dietrich, and H. M. Krause, “Anterior – posterior patterning in the Drosophila embryo,” vol. 12, 2002.

[31] D. Niessing, W. Driever, F. Sprenger, H. Taubert, H. Jäckle, and R. Rivera-Pomar, “Homeodomain position 54 specifies transcriptional versus translational control by Bicoid.,” Mol. Cell, vol. 5, no. 2, pp. 395–401, Feb. 2000, [Online]. Available: http://www.ncbi.nlm.nih.gov/pubmed/10882080.

[32] D. Niessing, S. Blanke, and H. Ja, “Bicoid associates with the 5 ؅ -cap-bound complex of caudal mRNA and represses translation,” pp. 2576–2582, 2002, doi: 10.1101/gad.240002.2576.

[33] H. G. Frohnhöfer and C. Nüsslein-Volhard, “Organization of anterior pattern in the Drosophila embryo by the maternal gene bicoid,” Nature, vol. 324, no. 6093, pp. 120–125, Nov. 1986, doi: 10.1038/324120a0.

[34] Q. Liu et al., “Ancient mechanisms for the evolution of the bicoid homeodomain’s function in fly development.,” Elife, vol. 7, Oct. 2018, doi: 10.7554/eLife.34594.

[35] M. J. Harms and J. W. Thornton, “Analyzing protein structure and function using ancestral gene reconstruction,” Current Opinion in Structural Biology, vol. 20. pp. 360–366, 2010, doi: 10.1016/j.sbi.2010.03.005.

[36] J. Treisman, P. Gönczy, M. Vashishtha, E. Harris, and C. Desplan, “A single amino acid can determine the DNA binding specificity of homeodomain proteins.,” Cell, vol. 59, no. 3, pp. 553–62, Nov. 1989, doi: 10.1016/0092-8674(89)90038-X.

[37] M. B. Noyes, R. G. Christensen, A. Wakabayashi, G. D. Stormo, M. H. Brodsky, and S. A. Wolfe, “Analysis of Homeodomain Specificities Allows the Family-wide Prediction of Preferred Recognition Sites,” Cell, 2008, doi: 10.1016/j.cell.2008.05.023.

[38] J. M. Baird-Titus, K. Clark-Baldwin, V. Dave, C. A. Caperelli, J. Ma, and M. Rance, “The solution structure of the native K50 Bicoid homeodomain bound to the consensus TAATCC DNA-binding site.,” J. Mol. Biol., vol. 356, no. 5, pp. 1137–51, Mar. 2006, doi: 10.1016/j.jmb.2005.12.007.

[39] R. Kraut and M. Levine, “Spatial regulation of the gap gene giant during Drosophila development,” Development, vol. 111, no. 2, 1991.

[40] E. D. Eldon and V. Pirrotta, “Interactions of the Drosophila gap gene giant with maternal and zygotic pattern-forming genes,” Development, vol. 111, no. 2, 1991.

[41] J. Mohler, E. D. Eldon, and V. Pirrotta, “A novel spatial transcription pattern associated with the segmentation gene, giant, of Drosophila.,” EMBO J., vol. 8, no. 5, pp. 1539–48, May 1989, Accessed: Jun. 28, 2020. [Online]. Available: http://www.ncbi.nlm.nih.gov/pubmed/2504582.

[42] M. Simpson-Brose, J. Treisman, and C. Desplan, “Synergy between the hunchback and bicoid morphogens is required for anterior patterning in Drosophila.,” Cell, vol. 78, no. 5, pp. 855–65, Sep. 1994, [Online]. Available: http://www.ncbi.nlm.nih.gov/pubmed/8087852.

[43] A. Porcher and N. Dostatni, “The Bicoid Morphogen System,” Curr. Biol., vol. 20, no. 5, pp. 249–254, 2010, doi: 10.1016/j.cub.2010.01.026.

[44] A. Ochoa-Espinosa et al., “The role of binding site cluster strength in Bicoid-dependent patterning in Drosophila.,” Proc. Natl. Acad. Sci. U. S. A., vol. 102, no. 14, pp. 4960–5, Apr. 2005, doi: 10.1073/pnas.0500373102.

[45] M. D. Schroeder, C. Greer, and U. Gaul, “How to make stripes: deciphering the transition from non-periodic to periodic patterns in Drosophila segmentation.,” Development, vol. 138, no. 14, pp. 3067–78, Jul. 2011, doi: 10.1242/dev.062141.

[46] G. Struhl, P. Johnston, and P. A. Lawrence, “Control of Drosophila body pattern by the hunchback morphogen gradient,” Cell, vol. 69, no. 2, pp. 237–249, Apr. 1992, doi: 10.1016/0092-8674(92)90405-2.

[47] X. Wu, V. Vasisht, D. Kosman, J. Reinitz, and S. Small, “Thoracic Patterning by the Drosophila Gap Gene hunchback,” Dev. Biol., vol. 237, no. 1, pp. 79–92, Sep. 2001, doi: 10.1006/DBIO.2001.0355.

[48] M. Hülskamp, C. Pfeifle, and D. Tautz, “A morphogenetic gradient of hunchback protein organizes the expression of the gap genes Krüppel and knirps in the early Drosophila embryo,” Nature, vol. 346, no. 6284, pp. 577–580, Aug. 1990, doi: 10.1038/346577a0.

[49] D. Yu and S. Small, “Precise registration of gene expression boundaries by a repressive morphogen in Drosophila.,” Curr. Biol., vol. 18, no. 12, pp. 868–76, Jun. 2008, doi: 10.1016/j.cub.2008.05.050.

[50] F. Liu, A. H. Morrison, and T. Gregor, “Dynamic interpretation of maternal inputs by the Drosophila segmentation gene network,” Proc. Natl. Acad. Sci., vol. 110, no. 17, pp. 6724–6729, Apr. 2013, doi: 10.1073/pnas.1220912110.

[51] Z. Xu, H. Chen, J. Ling, D. Yu, P. Struffi, and S. Small, “Impacts of the ubiquitous factor Zelda on Bicoid-dependent DNA binding and transcription in Drosophila.,” Genes Dev., vol. 28, no. 6, pp. 608–21, Mar. 2014, doi: 10.1101/gad.234534.113.

[52] M. Mir et al., “Dense Bicoid hubs accentuate binding along the morphogen gradient.,” Genes Dev., vol. 31, no. 17, pp. 1784–1794, Sep. 2017, doi: 10.1101/gad.305078.117.

[53] R. R. Datta et al., “A feed-forward relay integrates the regulatory activities of Bicoid and Orthodenticle via sequential binding to suboptimal sites.,” Genes Dev., vol. 32, no. 9–10, pp. 723–736, May 2018, doi: 10.1101/gad.311985.118.

[54] C. E. Hannon, S. A. Blythe, and E. F. Wieschaus, “Concentration dependent chromatin states induced by the bicoid morphogen gradient,” Elife, vol. 6, p. e28275, Sep. 2017, doi: 10.7554/eLife.28275.

[55] R. Schröder, “The genes orthodenticle and hunchback substitute for bicoid in the beetle Tribolium,” Nature, vol. 422, no. 6932, pp. 621–625, Apr. 2003, doi: 10.1038/nature01536.

[56] J. A. Lynch, A. E. Brent, D. S. Leaf, M. Anne Pultz, and C. Desplan, “Localized maternal orthodenticle patterns anterior and posterior in the long germ wasp Nasonia,” Nature, vol. 439, no. 7077, pp. 728–732, Feb. 2006, doi: 10.1038/nature04445.

[57] Q. Gao and R. Finkelstein, “Targeting gene expression to the head: the Drosophila orthodenticle gene is a direct target of the Bicoid morphogen,” Development, vol. 125, no. 21, pp. 4185–4193, Nov. 1998, Accessed: Jan. 12, 2016. [Online]. Available: http://dev.biologists.org/content/125/21/4185.abstract?ijkey=93cb26aef4cb3eef47761ecc2ee591bbc2345234&keytype2=tf_ipsecsha.

[58] J. Lynch and C. Desplan, “Evolution of development: beyond bicoid.,” Curr. Biol., vol. 13, no. 14, pp. R557–9, Jul. 2003, doi: 10.1016/s0960-9822(03)00472-x.

[59] Y. Yoon et al., “Embryo polarity in moth flies and mosquitoes relies on distinct old genes with localized transcript isoforms,” Elife, vol. 8, Oct. 2019, doi: 10.7554/eLife.46711.

[60] J. Klomp et al., “A cysteine-clamp gene drives embryo polarity in the midge Chironomus,” Science (80-.)., vol. 348, no. 6238, pp. 1040–1042, May 2015, doi: 10.1126/science.aaa7105.

[61] J. R. Bateman, A. M. Lee, and C. Wu, “Site-specific transformation of Drosophila via phiC31 integrase-mediated cassette exchange.,” Genetics, vol. 173, no. 2, pp. 769–77, Jun. 2006, doi: 10.1534/genetics.106.056945.

[62] S. Small, “In vivo analysis of lacZ fusion genes in transgenic Drosophila melanogaster,” Methods Enzymol., 2000, doi: 10.1016/s0076-6879(00)26052-7.

[63] D. Kosman, S. Small, and J. Reinitz, “Rapid preparation of a panel of polyclonal antibodies to Drosophila segmentation proteins,” Dev. Genes Evol., vol. 208, no. 5, pp. 290–294, 1998, doi: 10.1007/s004270050184.

[64] X. Wu, R. Vakani, and S. Small, “Two distinct mechanisms for differential positioning of gene expression borders involving the Drosophila gap protein giant,” Development, vol. 125, no. 19, 1998.

[65] L. E. Anderson, “Hoyer’s Solution as a Rapid Permanent Mounting Medium for Bryophytes,” Bryologist, vol. 57, no. 3, p. 242, Sep. 1954, doi: 10.2307/3240091.

[66] C. T. Rueden et al., “ImageJ2: ImageJ for the next generation of scientific image data,” BMC Bioinformatics, vol. 18, no. 1, p. 529, Dec. 2017, doi: 10.1186/s12859-017-1934-z.

[67] C. A. Schneider, W. S. Rasband, and K. W. Eliceiri, “NIH Image to ImageJ: 25 years of image analysis,” Nat. Methods, vol. 9, no. 7, pp. 671–675, Jul. 2012, doi: 10.1038/nmeth.2089.

