## Supplemental Figures and legends for "Suboptimal intermediates underlie evolution of the Bicoid homeodomain"

#### Supplementary Figures S1-S3

Pinar Onal, Himari Imaya Gunasinghe, Kristaley Yui Umezawa, Michael Zheng, Jia Ling, Tasmima Tazin, Leen Azeez, Anecine Dalmeus, Stephen Small*

New York University, Department of Biology

### Figure S1

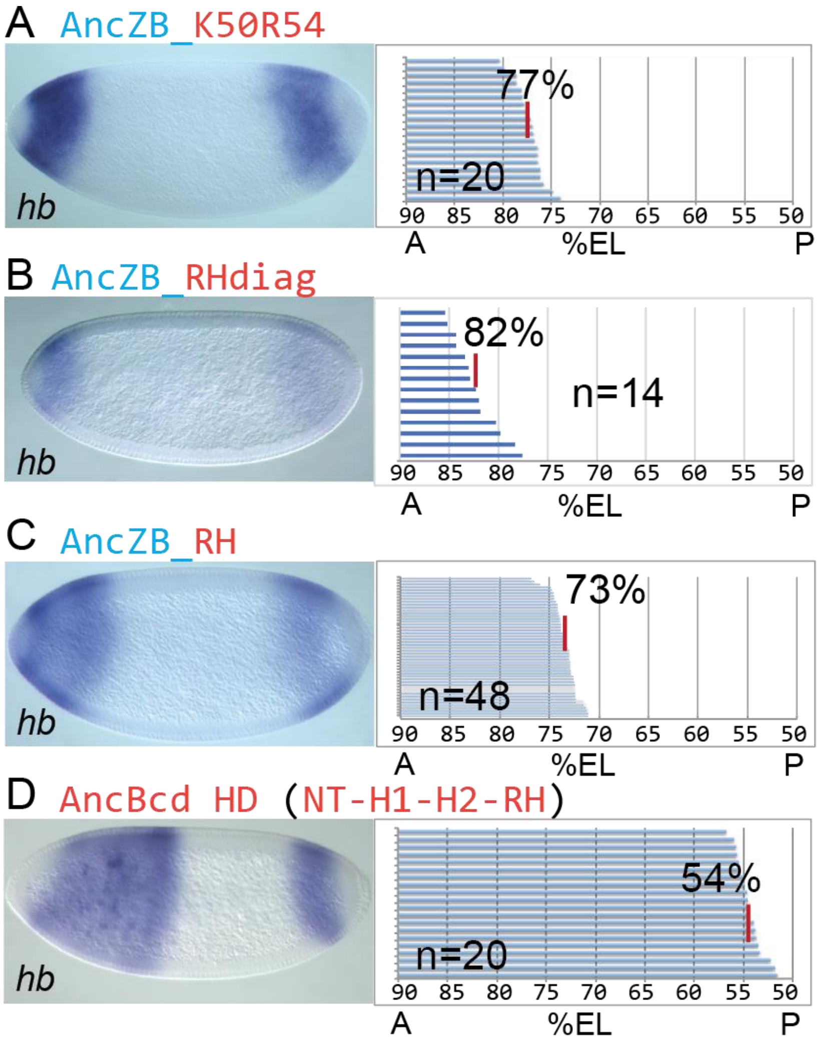


*Figure S1: hb expression patterns and hb posterior boundary positions (pbps) of anterior hb patterns of stage 5 embryos from transgenic lines described in Figure 2 and control AncBcd HD lines. Related to Figure 2.* A-D. Left panels show representative stage 5 embryos stained by *in situ* hybridization to detect *hb*. Same embryos as in Figure 2 are chosen as representative embryos. Panels to the right show the analysis of multiple embryos to calculate the average pbp (% EL; anterior tip = 100%) of *hb* at stage 5. Each horizontal line in each panel represents the anterior *hb* expression pattern in a single embryo, and the average pbp is denoted by a vertical red line. N numbers of embryos are used for each experiment. The most anterior tip is 100% EL (embryo length). Only 90 -50 % EL is shown on the left panels.

### Figure S2

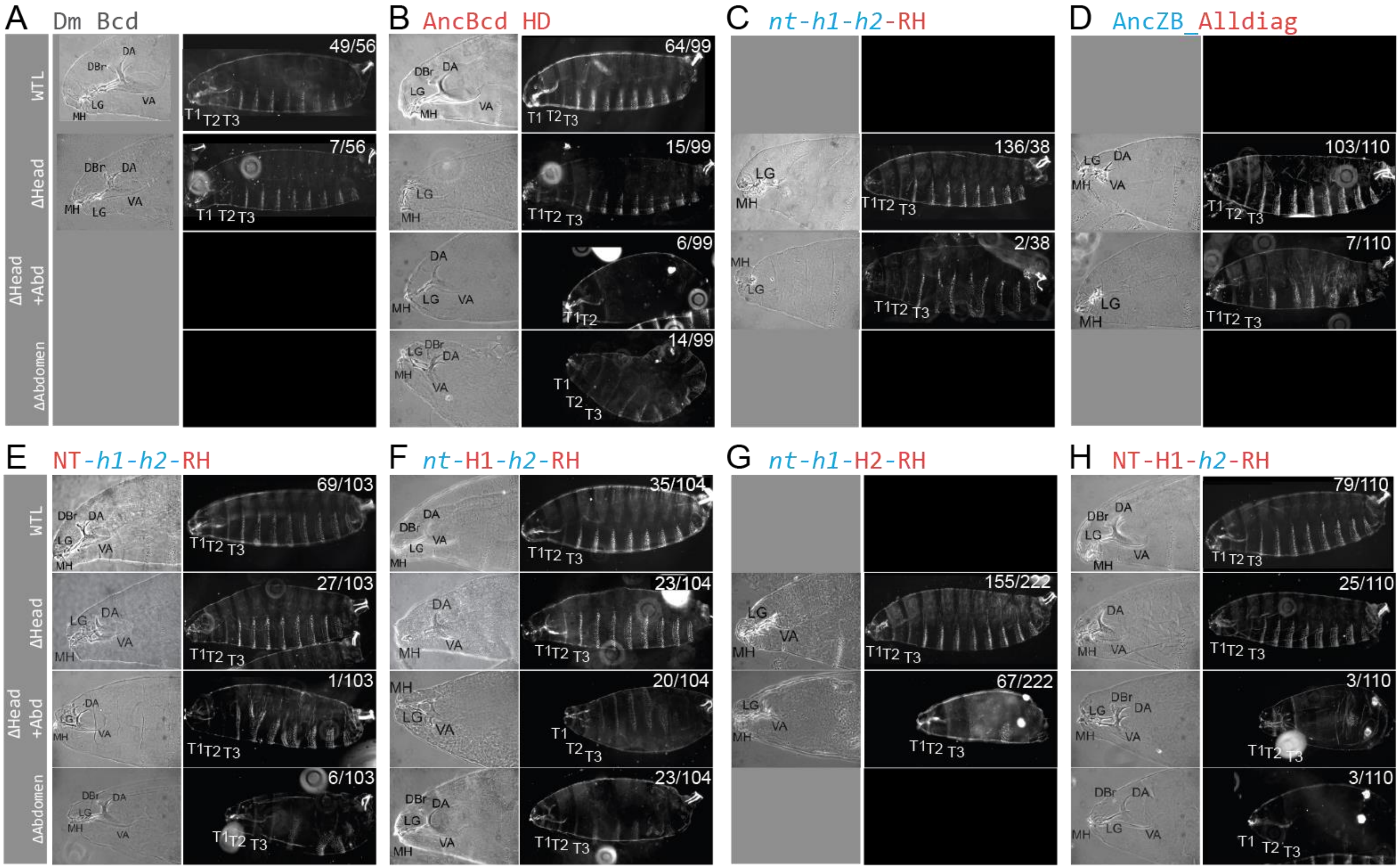


Figure S2: Phenotypic variation in transgenic D.melanogaster expressing ancestral proteins. Related to Figures 3 and 4. **A-H**. Representative of first instar larvae showing different degrees of morphological rescue upon expression of different chimeric ancestral HD. Morphologies of individual larvae were classified as described in the text and Methods. Left of each panel shows cuticle preparations of the head, and the right shows cuticle of the whole body of the transgenic larvae. Ratio given is the ratio of number of larvae counted for the category per the total larvae in one transgenic line.

### Figure S3

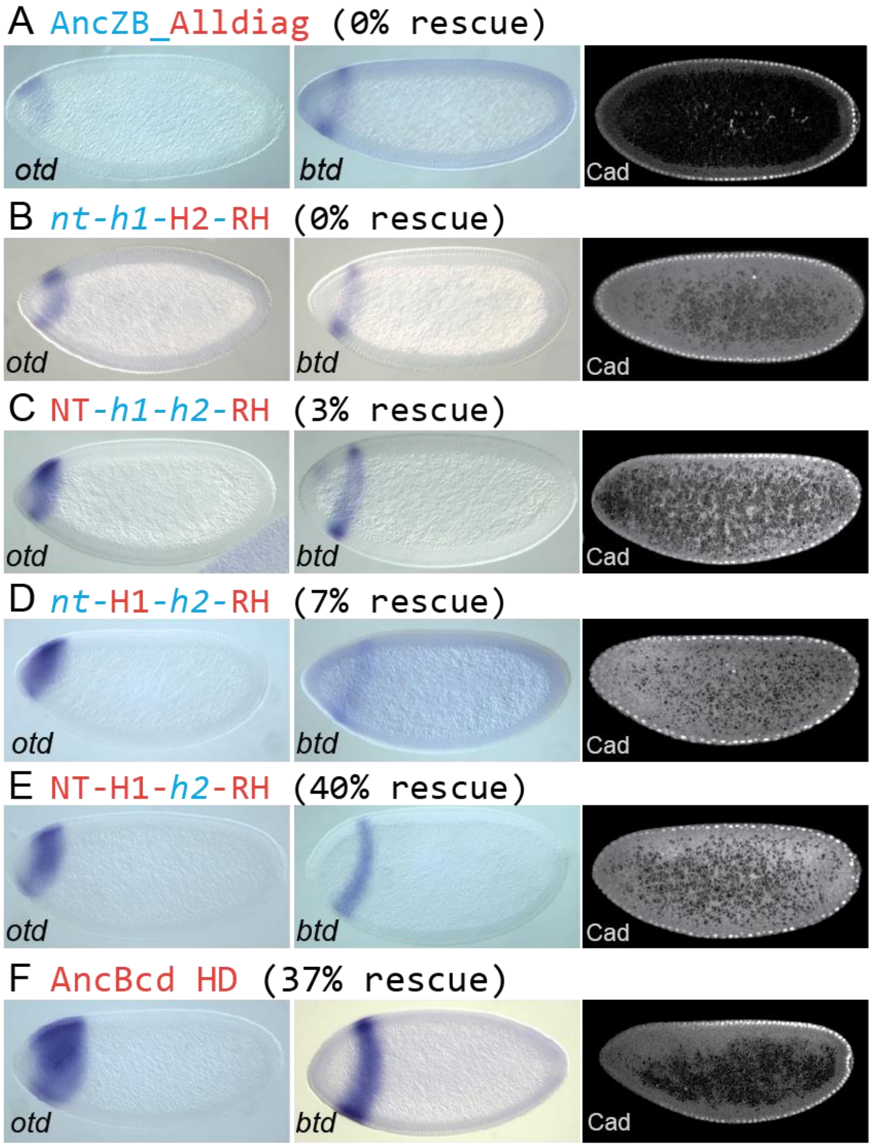


*Figure S3: Transcriptional and translational activities of constructs that were described in figures 3 and 4. Related to Figures 3 and 4.* A-F. RNA expression of head gap genes *otd (orthodenticle)* and *btd (button head);* and Caudal (Cad) protein distribution.
